# Protection and deprotection of the Rec8 cohesin complex during meiosis

**DOI:** 10.1101/2025.09.19.677360

**Authors:** Yongxin Liu, Ke Zhang, Ziteng Bai, Li Sun, Haitong Hou, Yoshinori Watanabe

## Abstract

In the first meiotic division, the cohesin Rec8 is cleaved by separase along the chromosome arms but is protected at the centromere by shugoshin (Sgo1), maintaining centromeric cohesion throughout meiosis I. Another meiotic regulator, meikin (Moa1), supports the protective function of Sgo1 primarily by phosphorylating S450 of Rec8. Both Sgo1 and Moa1 are meiosis I-specific proteins. Unlike fission yeast Sgo1, mammalian shugoshin (SGO2) persists at centromeres throughout meiosis I and II, but REC8 cohesin is protected only during meiosis I. Here, we show that fission yeast Moa1 and Sgo1 appear at kinetochores/centromeres from prophase I to metaphase I and are degraded during anaphase I by the APC/C-Slp1 pathway. We expressed non-degradable versions of Moa1 and Sgo1 in meiosis II and investigated how these proteins protect the cohesin Rec8 during meiosis II. Our analyses suggest that the localization of Sgo1 and the phosphorylation of Rec8 at S449 and S450 are necessary and sufficient events for protecting Rec8 cohesin during meiosis II. The absence of either event leads to deprotection at meiosis II.

## Introduction

The cohesin complex, which mediates sister chromatid cohesion and DNA looping, plays a crucial role in chromosome compaction and separation during mitosis (Yatskevich et al. 2019; Davidson and Peters 2021). Cohesin comprises two SMC (structural maintenance of chromosome) family proteins, which in fission yeast (*S. pombe*) are, called Psm1 and Psm3, a kleisin subunit (*S. p*. Rad21), and an accessory subunit (*S. p*. Psc3) (Tomonaga et al. 2000). In proliferating cells, sister chromatid cohesion is established during S phase and maintained until metaphase when the sister chromatids are captured by spindle microtubules from opposite poles. To initiate anaphase, the anaphase-promoting complex (APC/C) triggers the degradation of securin, an inhibitory chaperone for separase, a protease. Separase then cleaves Rad21 along the entire chromosome. This triggers the separation of sister chromatids and their movement to opposite poles, a process called equational division (Yanagida 2000; Nasmyth 2001). In meiosis, the cohesin subunit Rad21 is largely replaced by the meiosis-specific Rec8 protein. This ‘cohesin switch’ enables reductional, rather than equational, division during meiosis I (Watanabe and Nurse 1999; Watanabe et al. 2001). During the meiosis I division, sister kinetochores attach to microtubules emanating from the same pole (mono-orientation). Rec8 is cleaved only along the chromosome arms but not at the centromere because the protein shugoshin (Sgo1) protects Rec8 from cleavage by separase at the centromere (cohesion protection) (Kitajima et al. 2004; Marston 2014). PP2A bound to Sgo1 antagonizes casein kinase 1 (CK1)-dependent Rec8 phosphorylation.

Because this phosphorylation is a prerequisite for cleavage by separase, Sgo1 protects cohesion at the centromeres (Kitajima et al. 2006; Riedel et al. 2006; Ishiguro et al. 2010; Katis et al. 2010). The meiosis-specific kinetochore regulator meikin (*S*.*p*. Moa1) bound to Plo1 kinase also contributes to cohesion protection in addition to mono-orientation (Shonn et al. 2002; Katis et al. 2004; Lee et al. 2004; Yokobayashi and Watanabe 2005; Kim et al. 2015; Miyazaki et al. 2017). The phosphorylation of Rec8-S450 by Moa1-Plo1 activates the Sgo1-PP2A complexes, thereby promoting cohesion protection at the centromere (Ma et al. 2021). Because Sgo1 is degraded in anaphase I and absent in meiosis II, it is reasonable to hypothesize that Sgo1 degradation is the reason for ‘deprotection’ of Rec8 cohesin in meiosis II in fission yeast. However, in mammals, SGO2 (the Sgo1 homolog) localizes to centromeres not only in meiosis I but also in meiosis II, suggesting that meiosis II specific deprotection mechanisms exist. Previous studies showed that SGO2 colocalizes with Rec8 cohesin at the inner centromere during metaphase I when sister kinetochores are attached to the same pole and there is no tension between sister kinetochores. However, in metaphase II, when sister kinetochores are pulled toward opposite poles, SGO2 separates from the Rec8 cohesin site and moves to the inner face of the kinetochore (Gomez et al. 2007; Lee et al. 2008). These observations led us to hypothesize that tension between sister kinetochores may contribute to the deprotection of Rec8 cohesin during meiosis II. Another study suggested that colocalization of SGO2-PP2A with the PP2A inhibitor SET causes deprotection in mammalian oocytes (Chambon et al. 2013). In budding yeast, though, Sgo1 also localizes to centromeres during meiosis II, as in mammals, yet, deletion of the SET homologs does not interfere with chromosome segregation in meiosis II (Jonak et al. 2017). This suggests that deprotection by SET is not widely conserved and that other deprotection mechanisms must exist. Because co-expression of Sgo1 and Mps1 at meiosis II protects Rec8 in budding yeast, it was suggested that the APC/C-dependent degradation of Sgo1 and Mps1 kinase during anaphase I may be the reason for deprotection in meiosis II (Jonak et al. 2017; Mengoli et al. 2021). However, although Mps1 is a part of the spindle assembly checkpoint (SAC) and can delay anaphase onset, its substrate in promoting protection has not been uncovered. Thus, it is unknown exactly which regulation allows for deprotection and ectopic protection in meiosis II in budding yeast. To address this long-standing ‘deprotection’ problem, we reconstituted ectopic protection in meiosis II of fission yeast and determined which exact proteins and regulations are missing in physiological meiosis II.

## Results and Discussion

### Sgo1 is degraded in anaphase I in a D-box dependent manner

Because fission yeast Sgo1 is degraded at anaphase I in an APC/C-Slp1-dependent manner (Kitajima et al. 2004), we searched the Sgo1 protein sequence for a putative D-box sequence, and identified <u>K</u>CI<u>L</u>KDKT<u>N</u> at residues 182-190 of Sgo1 (Fig. 1A). To determine whether the motif is important for Sgo1 degradation, we introduced a <u>D</u>-<u>b</u>ox mutation L185A in the endogenous *sgo1* gene (*sgo1-DB*) and examined protein expression during meiosis (Fig. 1B). Wild-type Sgo1 appeared at centromeres from prophase I to metaphase I and disappeared at anaphase I. In contrast, Sgo1-DB localized at the centromere not only in metaphase I but also in metaphase II (Fig. 1C). To determine the consequence on chromosome segregation, we used a *lacO* array integrated at the centromere of chromosome I and visualized by LacI-GFP (*imr1*::GFP) (Fig. 1D). Remarkably, meiotic chromosome segregation was largely intact in *sgo1-DB* cells (Fig. 1E), indicating that the timely degradation of Sgo1 at anaphase I and its absence in meiosis II is not important for proper meiotic chromosome segregation. These results support the notion that, similar to budding yeast and mouse oocytes, shugoshin localization per se is not sufficient to protect Rec8 during meiosis II in fission yeast.

**Figure 1.**
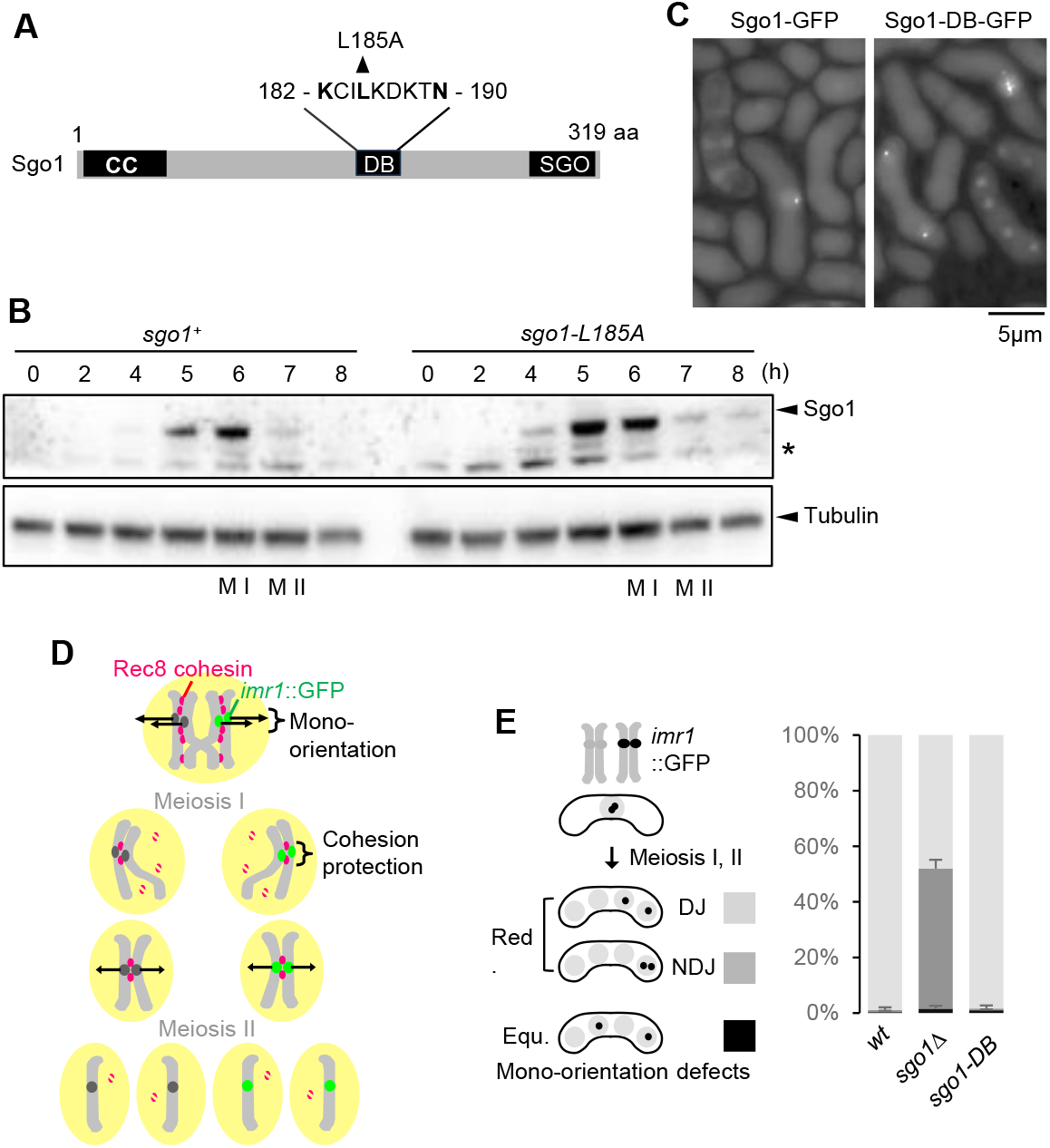
Sgo1 is degraded in anaphase I by the APC/C-Slp1 pathway. **A**, Schematic diagram of the Sgo1 protein showing the PP2A-interacting coiled-coil region (CC), D-box (DB), and SGO motif (SGO). D-box sequence and mutation site are shown. **B**, *sgo1-GFP* and *sgo1-DB-GFP* cells were induced to undergo synchronous meiosis by *pat1-114* inactivation. Immunoblot analysis was performed with anti-GFP and anti-tubulin antibodies by using lysates prepared from the indicated cells at each time point. The asterisk indicates a nonspecific band. M I: meiosis I, M II: meiosis II. **C**, *sgo1-GFP* and *sgo1-DB-GFP* cells were spotted on an SPA plate and examined for the localization of Sgo1 during meiosis. **D**, Schematic depiction of behaviors of homologous chromosomes and Rec8 cohesin during meiosis, showing *imr1*::GFP marked on one homolog. **E**, Segregation pattern of *imr1*::GFP marked on one homolog in the indicated cells was monitored in cells after the meiosis II division. DJ: disjunction, meaning proper sister chromatid segregation in meiosis II. NDJ: nondisjunction, meaning sister chromatids move to the same pole in meiosis II. Red. (reductional): sister chromatids moved to the same pole in meiosis I. Equ. (equational): sister chromatids moved to opposite poles already in meiosis I. Error bars represent SD (3 independent experiments, n > 150 cells in each experiment).

### Sgo1-DB protects Rec8-2E (S449E, S450E) in meiosis II

Our previous studies showed that phosphorylation of Rec8-S412 is required for cleavage of Rec8 (at R384) by separase (Ishiguro et al. 2010). In contrast, phosphorylation of Rec8 at S449 and S450 is required to protect Rec8 from cleavage. The corollary is that separase recognizes the phosphorylation at S412, thereby promoting cleavage of Rec8. Conversely, Sgo1-PP2A recognizes the phosphorylation at S449 and S450 of Rec8 and dephosphorylates pS412 to prevent separase-mediated cleavage (Fig. 2A). Accordingly, *rec8-2A* (non-phosphorylatable mutant *rec8-S449A, S450A*) impairs the protection of Rec8, whereas *rec8-2E* (phosphomimetic mutant *rec8-S449E, S450E*) enhances the protection (Ma et al. 2021). Consistent with our previous study, in *rec8-2E* cells arrested at prometaphase II by the *mes1-B44* mutation, homologous chromosomes (homologs) connected by chiasmata did not separate (non-disjunction; NDJ) in >70% of cells and this NDJ defect was suppressed by *sgo1Δ* (Fig. 2B). This suggests that Sgo1 protected Rec8-2E not only at the centromeres but also along chromosome arms, although the amount of Sgo1 on the arms was low (Fig. 2B). We next addressed whether Rec8-2E is protected during meiosis II in *rec8-2E* cells expressing mCherry-Atb2 (α2-tubulin). Microscopic observation indicated that nuclear division was blocked in meiosis I of *rec8-2E* cells, despite proper spindles formation (Fig. 2C), consistent with the chromosome NDJ observed (Fig. 2B). However, in meiosis II, nuclear division occurred, although the two spindles were irregularly formed in the single nucleus (Fig. 2C). Accordingly, in *rec8-2E* cells, single chromatid marked by *imr1*::GFP distributed into the forming spores, albeit with some degree of mis-segregation (Fig. 2D). These results suggest that Rec8-2E was protected during meiosis I depending on Sgo1, but not in meiosis II most likely because Sgo1 is absent in meiosis II (Fig. 1C). To address whether Sgo1 can protect Rec8-2E in meiosis II, we examined the meiotic chromosome segregation in *sgo1-DB rec8-2E* cells, in which Sgo1 protein localizes at centromeres in both meiosis I and II (Fig. 1C). These cells produced one or two spores and most zygotes showed *imr1*::GFP signals within one nucleus (Fig. 2D). These results suggest that neither sisters nor homologs separated in both meiosis I and II in *sgo1-DB rec8-2E* cells. Hence, deprotection of *rec8-2E* in meiosis II requires Sgo1 degradation (Fig. 2E).

**Figure 2.**
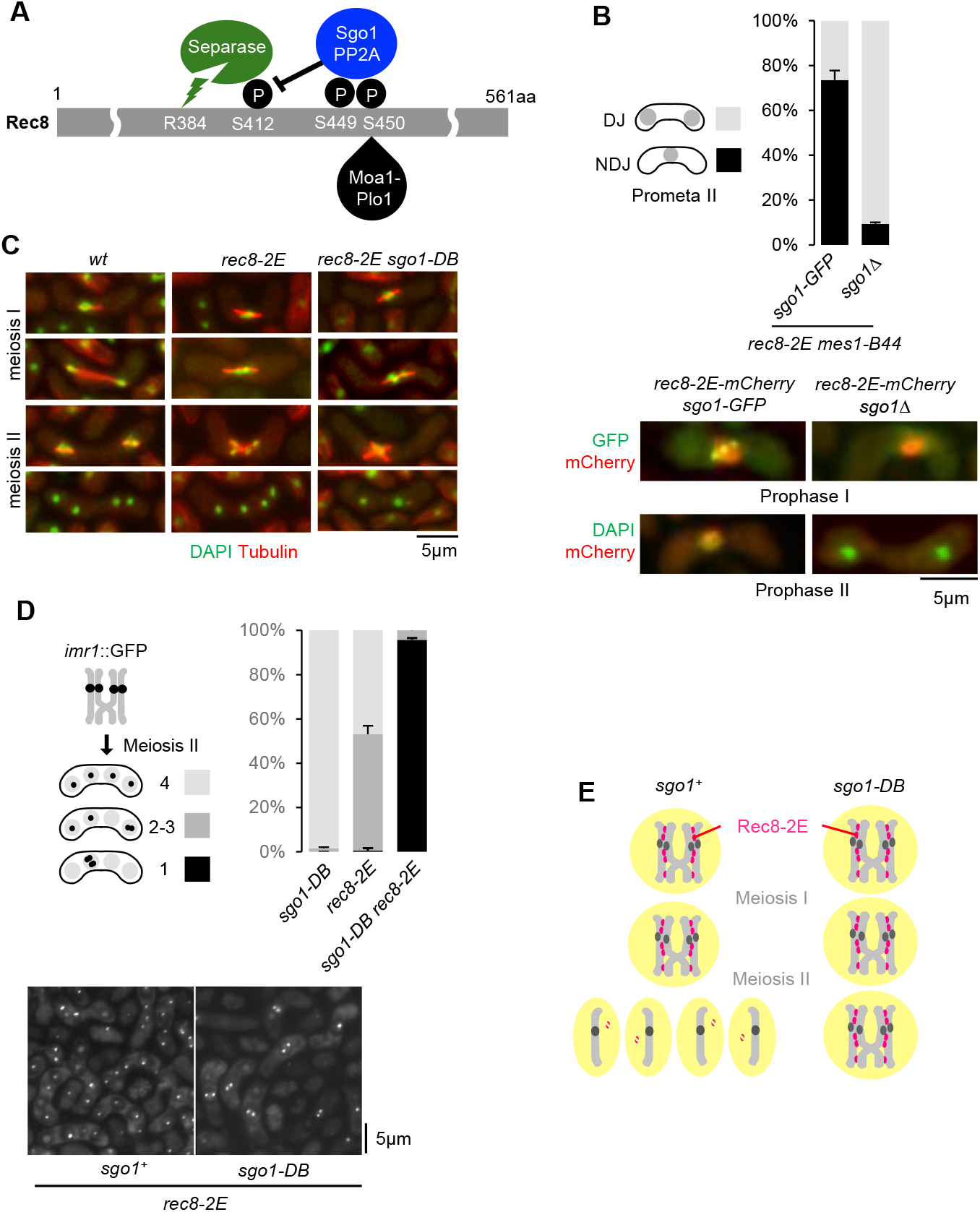
Sgo1-DB protects Rec8-2E (S449E, S450E) in meiosis II. **A**, Schematic diagram of how Rec8 phosphorylation and Sgo1-PP2A localization regulate Rec8 cleavage by separase. Phosphorylation at S412 of Rec8 mediated by CK1 is required for separase-mediated Rec8 cleavage at R384, whereas both phosphorylation at S449 and S450 of Rec8 are required for the protective function of Sgo1-PP2A. **B**, The indicated cells arrested at prometaphase II by *mes1-B44* were fixed and stained with DAPI to monitor meiotic nuclear divisions. Representative images of the indicated cells at prophase I and prophase II are shown. **C**, The indicated cells are shown undergoing meiosis I and meiosis II by staining tubulin and DNA. **D**, Segregation pattern of *imr1*::GFP marked on both homologs was monitored in the indicated cells after the meiosis II division. Representative images of the indicated cells are shown. **E**, Schematic diagram of behaviors of homologous chromosomes and Rec8 cohesin during meiotic divisions of *rec8-2E sgo1*^*+*^ and *rec8-2E sgo1-DB* cells.

### Sgo1-DB protects Rec8-2E at centromeres in *rec11Δ* cells with reduced arm cohesion

In meiosis, Rec8 forms a complex with the Psc3 subunit mostly around the centromeres and with the meiosis-specific Rec11 subunit along the chromosome arms. Therefore, in *rec11Δ* cells, cohesion is diminished along the chromosome arms but preserved at centromeres due to the Rec8-Psc3 complexes (Kitajima et al. 2003). Thus, we introduced the *rec11Δ* mutation in *rec8-2E* cells to reduce the effects of persistent arm cohesion. To delineate the protection of centromeric Rec8-2E in meiosis II, the dynamics of *imr1::*GFP on one homolog were observed in *rec8-2E rec11Δ* cells by time-lapse imaging. Simultaneously, DNA was visualized by Pht1 (histone H2A variant, H2A.Z) fused with mCherry. During anaphase I, DNA was stretched out but subsequently became round, implying that homologs were not completely separated in anaphase I (Fig. 3A, *sgo1*^*+*^), possibly due to the residual cohesion and protection along the arms even in the absence of Rec11. In the following meiosis II, DNA were separated into four globules, and most sister *imr1::*GFP were separated during meiosis II, indicative of the absence of protection (Fig. 3B,C, *sgo1*^*+*^). In contrast, in *rec8-2E rec11Δ sgo1-DB* cells, despite showing transient splitting of *imr1::*GFP indicative of bi-orientation at anaphase II, sister chromatids did not separate (Fig. 3A,B,C, *sgo1-DB*). When the mutant Sgo1-N29I-DB, which abolishes PP2A binding, was expressed instead of Sgo1-DB, the protection was largely lost, indicating that protection is indeed dependent on PP2A bound to Sgo1 (Fig. EV1).

**Figure 3.**
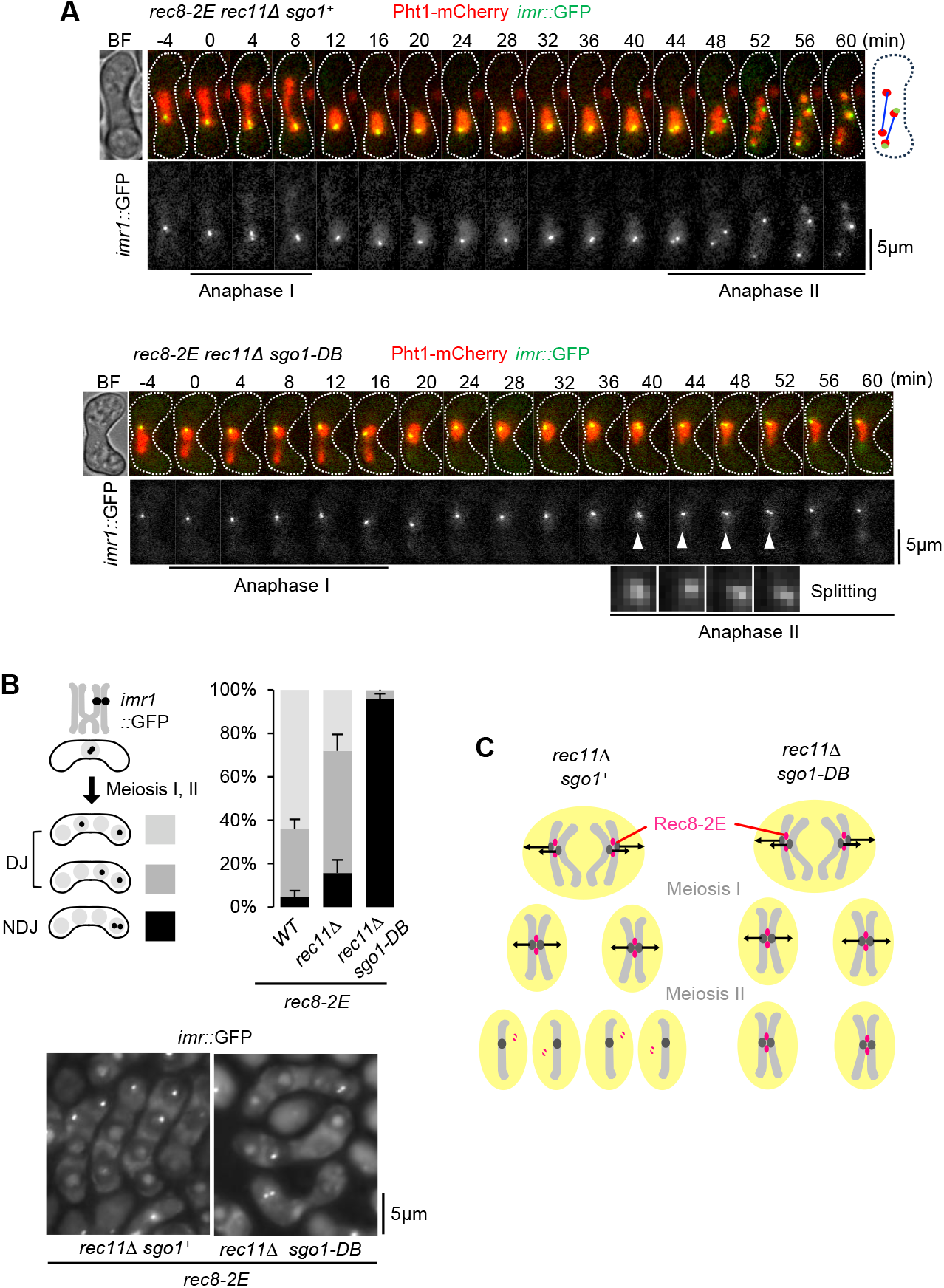
Sgo1-DB protects Rec8-2E at centromeres in *rec11Δ* cells with reduced arm cohesion. **A**, Sister centromere behavior was observed in the indicated strains having *imr1*::GFP on only one homolog. DNA was visualized by expressing Pht1-mCherry from the endogenous locus. The representative examples of the indicated strains are shown from the beginning of the first anaphase to the end of the second anaphase. BF, bright field. Anaphase I and II are indicated by lines. **B**, Segregation pattern of *imr1*::GFP marked on one homolog was monitored in the indicated cells after the meiosis II division. Note that in *rec8-2E sgo1-DB* cells, sister chromatids entirely move to the same pole in meiosis II, indicative of ectopic cohesion protection during anaphase II. Representative images of the indicated cells are shown. **C**, Schematic depiction of behaviors of homologous chromosomes and Rec8 cohesin during meiotic divisions of *rec11Δ rec8-2E sgo1*^*+*^ and *rec11Δ rec8-2E sgo1-DB* cells.

### Sgo1-DB and Moa1-DB protect Rec8-S449E in meiosis II

Given that Sgo1 can protect Rec8-2E in meiosis II, we next examined whether Sgo1 can protect Rec8-S449E or Rec8-S450E (only one phosphomimetic mutation). Previously, we showed that Plo1 bound to Moa1 phosphorylates at S450 of Rec8, thereby contributing to the protection of Rec8 during meiosis I. We therefore explored whether Moa1 can also contribute to the protection of Rec8 in meiosis II. Notably, like Sgo1, Moa1 also carries a D-box sequence <u>R</u>PA<u>L</u>QDKT<u>N</u> (residues 32-40) (Zhou et al. 2025) (Fig. 4A). We introduced the <u>D</u>-<u>b</u>ox mutation L35A into endogenous Moa1 (Moa1-DB) and examined its protein expression and localization during meiosis (Fig. 4B,C). While wild-type Moa1 appeared at kinetochores from prophase I to metaphase I and disappeared in anaphase I, Moa1-DB persisted at centromeres throughout meiosis even after anaphase II (Fig. 4C). In *moa1-DB* cells, meiotic chromosome segregation was largely intact (Fig. 4D), indicating that the timely degradation of Moa1 at anaphase I or its absence in meiosis II is not important for proper meiotic chromosome segregation in fission yeast. Rec8-S449E was partially protected by Sgo1-DB (39% NDJ), whereas Rec8-S450E was not protected (3% NDJ), similar to Rec8-WT (Fig. 4E). Expression of Moa1-DB increased the protection in *rec8-S449E sgo1-DB rec11Δ* cells from 39% to 78% (Fig. 4E). In contrast, expression of Moa1-DB did not increase the protection in *rec8-S450E sgo1-DB rec11Δ* cells, supporting the notion that Plo1 bound to Moa1-DB indeed phosphorylates S450, but not S449 or other sites of Rec8 to enhance the protective capacity of Sgo1. To date, the kinase that phosphorylates S449 of Rec8 remains unknown (Ma et al. 2021). However, our analyses suggest that Sgo1 localization and phosphorylation of Rec8 at S449 and S450 are required and sufficient events to protect Rec8 cohesin at the centromere in meiosis II, implying that the absence of either event causes ‘deprotection’ in meiosis II.

**Figure 4.**
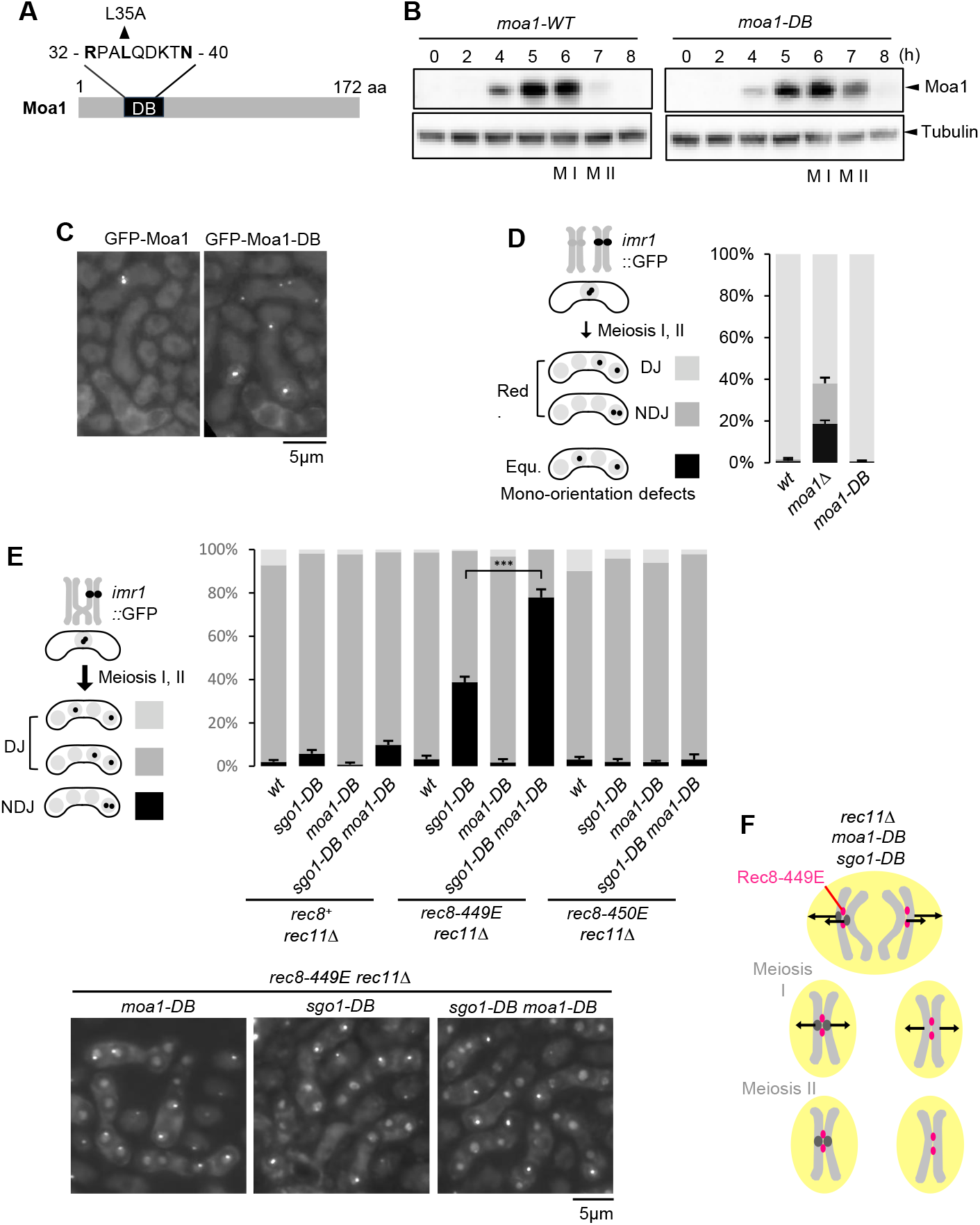
Sgo1-DB and Moa1-DB protect Rec8-S449E in meiosis II. **A**, Schematic diagram of the Moa1 protein showing the D-box (DB). D-box sequence and mutation site are shown. **B**, *GFP-PK-moa1*^*+*^ and *GFP-PK-moa1-DB* cells were induced to undergo synchronous meiosis by *pat1-114* inactivation. M I: meiosis I, M II: meiosis II. Immunoblot analysis was performed with anti-PK and anti-tubulin antibodies by using lysates prepared from the indicated cells at each time point. **C**, *GFP-PK-moa1*^*+*^ cells and *GFP-PK-moa1-DB-GFP* cells were spotted on an SPA plate and examined for the localization during meiosis. **D**, Segregation pattern of *imr1*::GFP marked on one homolog was monitored in the indicated cells after the meiosis II division. **E**, Segregation pattern of *imr1*::GFP marked on one homolog was monitored in the indicated cells after the meiosis II division. Representative images of the indicated cells are shown. **F**, Schematic diagram of behaviors of homologous chromosomes and Rec8 cohesin during meiotic divisions of the *rec11Δ rec8-449E sgo1-DB moa1-DB* cell.

### Moa1-bound Plo1 can be functionally replaced by Mps1

Our studies reveal that expression of Sgo1 and Rec8-2E (S449E, S450E) is sufficient for protection at meiosis II and that phosphorylation at S450 of Rec8 is primarily catalyzed by Moa1-bound Plo1. On the other hand, prior studies with budding yeast have shown that ectopic expression of Sgo1 and Mps1 kinase during meiosis II is sufficient for establishing protection of Rec8 (Mengoli et al. 2021). A study in mice oocytes suggested that MPS1 plays an important role in cohesion protection in meiosis I (El Yakoubi et al. 2017). Given that Plo1 (PLK1) and Mph1 (MPS1 pombe homolog) share similar phosphorylation consensus sites in mammals and fission yeast (von Schubert et al. 2015; Miyazaki et al. 2017), MPS1 and PLK1 may be functionally homologous. Rec8/cohesin, Sgo1/shugoshin and Moa1/meikin are widely conserved among eukaryotes, but no homologs of Moa1/meikin have been identified in plants so far (Singh et al. 2025), although the Rec8 cohesin-dependent mono-orientation mechanism is conserved (Chelysheva et al. 2005). Indeed, PLK1, the partner of Meikin, is missing in plants and is likely functionally replaced by MPS1 (de Carcer et al. 2011). To probe for the exchangeability of these kinases and explore potential evolutionary trajectories, we examined whether Moa1-bound Plo1 can be replaced by Mph1 in fission yeast meiosis. We used *rec12Δ mes1-B44* cells to test the Moa1-bound Plo1 function. In the absence of Rec12, homologs are not connected, and mono-orientation and protection defects in meiosis I will manifest as equational segregation in meiosis I, which we monitor in the *mes1-B44* prometaphase II arrest. Of note, both mono-orientation and protection are defective in *moa1-T101A* cells because Moa1-T101A fails to recruit Plo1 (Miyazaki et al. 2017). By fusing Plo1 to the N-terminus of Moa1-T101A, we tested whether fused Plo1 can restore Moa1 function at the kinetochore. Indeed, the mono-orientation defect in *moa1-T101A* cells (93% equational) was significantly suppressed in *plo1-moa1-T101A* cells (37% equational) and, also considerably, even though to a lesser extent, in *mph1-moa1-T101A* cells (54% equational) (Fig. 5A,B). Thus, Moa1-tethered Mph1 substituted for nearly 70% of the function of Moa1-tethered Plo1. Taken together, these results support the notion that PLK1 and MPS1 share important substrates and mechanisms required for meiotic chromosome segregation in yeasts and mammals.

**Figure 5.**
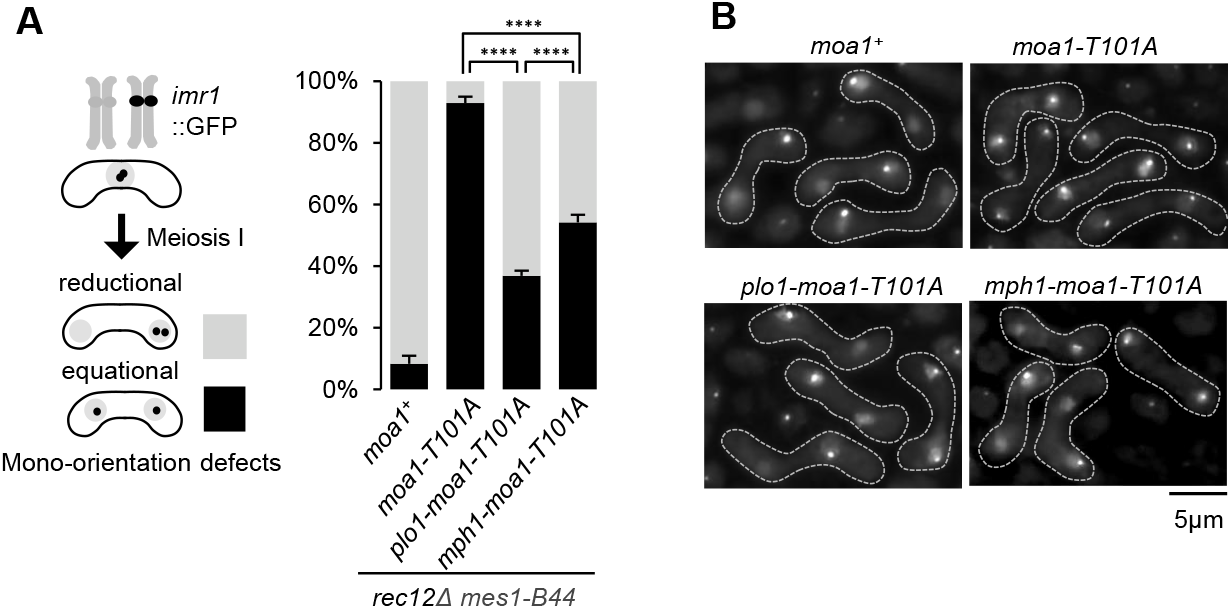
Moa1 bound Plo1 can be partially replaced by Mps1. **A**, The chromosome segregation pattern during meiosis I was counted by observing heterozygous *imr1*::GFP signals in prophase II arrest by *mes1-B44* in the indicated cells. Error bars, SD from three independent experiments. n > 150 cells in each experiment. **B**, Representative images of the indicated cells are shown.

### A mono-orientation-dependent protection model

A yeast two-hybrid assay revealed that Par1, the PP2A subunit which binds Sgo1, binds Rec8-2E but not Rec8-WT or Rec8-2A (Fig. EV2) (Ma et al. 2021). These results suggest that the dynamic interaction between the Sgo1-PP2A complex and Rec8 at the centromere depends on phosphorylation of Rec8 at S449 and S450. This is reminiscent of the binding of human Sgo1-PP2A complexes to cohesin, which is also regulated by phosphorylation and is essential for protecting cohesin at centromeres during mitosis (Liu et al. 2013). Importantly, during fission yeast meiosis, the phosphorylation of Rec8 is mediated in part by the kinetochore-localized Moa1-bound Plo1, which is geometrically closer to Rec8 in mono-oriented kinetochores than in bi-oriented kinetochores (Fig. 6A,B). Moreover, the microtubule pulling force at bi-oriented kinetochores generates tension, which increases the distance between kinetochore-associated kinases and Rec8, potentially reducing the phosphorylation of Rec8 required for its interaction with Sgo1 (Fig. 6B). On the other hand, the kinetochore-bound Bub1 kinase plays an important role in concentrating Sgo1 by phosphorylating H2A (Kawashima 2010). Thus, Bub1 contributes to the colocalization of Sgo1 and Rec8 at mono-oriented kinetochores during meiosis I, whereas at bi-oriented kinetochores during meiosis II, Bub1 may keep Sgo1 away from Rec8 while Rec8 is not efficiently phosphorylated by Plo1 or Mps1 (Fig. 6B). Rec8-2E may maintain the binding to Sgo1 even at bi-oriented kinetochores (Fig. 6C), preserving the protection during meiosis II as shown experimentally (Fig. 3A, *sgo1-DB*). Taken together, we propose a mono-orientation-dependent protection model in which phosphorylation-dependent interaction between Rec8 and Sgo1 is prominent at mono-oriented but less so at bi-oriented kinetochores (Fig. 6A,B), explaining why the ectopic presence of Sgo1 in meiosis II does not impair segregation. This model complements the tension-dependent deprotection model (Gomez et al. 2007; Lee et al. 2008) as well as the kinetochore individualization-dependent deprotection model (Gryaznova et al. 2021; Ogushi et al. 2021), which were both proposed in studies of mammalian oocytes.

**Figure 6.**
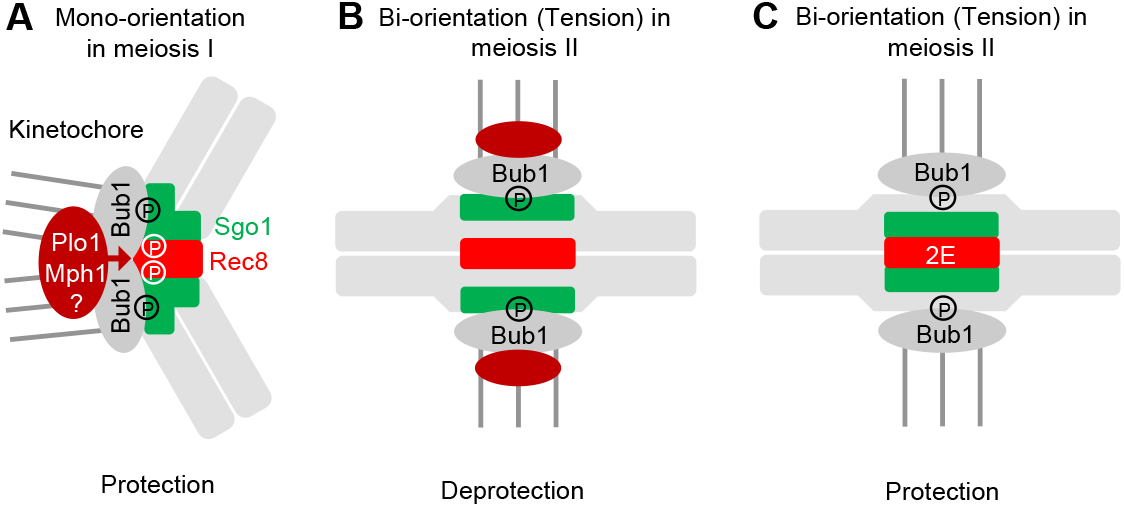
A mono-orientation-dependent protection model. **A**, Schematic diagram of mono-orientation-dependent protection model. **B**, Bi-orientation-dependent deprotection model. **C**, Rec8-2E is resistant to bi-orientation-dependent deprotection. See the text for further explanation.

## Materials and methods

### *Schizosaccharomyces pombe* strains and media

Unless otherwise stated, all media and growth conditions were as described previously (Moreno et al. 1991). *S. pombe* strains used in this study are described in Supplemental table S1. Complete medium (YE), minimal medium (SD and EMM) and sporulation media (SPA) were used. Deletion and tagging of *genes* by GFP or mCherry were carried out according to the PCR-based gene targeting method for *S. pombe* using the *kanMX6 (kanR), hphMX6 (hygR), bsdMX6(bsdR), natMX6 (natR*) and *aurMX6(aurR)* (Hashida-Okado et al. 1998) genes as selection markers (Bahler et al. 1998).

### Synchronous cultures of fission yeast meiotic cells

For meiosis microscopic observation, logarithmically growing cells were collected and resuspended in 20 mg/mL Leucine, spotted on SPA and incubated at 28°C. For chromosome segregation assay, *imr1*::GFP was observed in the *mes1*^*+*^ cells or the *mes1-B44* mutant that arrests at prophase II (after 16 h). For nuclei counting, DNA was visualized by staining with 2 μg/mL DAPI after fixing cells with 70% ethanol.

### Fluorescence microscopy

All fluorescence microscopy was performed using a Nikon ECLIPSE Ti2-E inverted microscope with photometrics PRIME 95B camera. This microscope was controlled by NIS-Elements software. Nine z sections (spaced by 0.4 μm each) of the fluorescent images were converted into a single two-dimensional image by Extended Depth of Focued (EDF) module. Image J software (NIH) was used to adjust brightness and contrast.

### Time-lapse imaging

Live-cell recordings were performed at 25°C using a Nikon ECLIPSE Ti2-E inverted microscope with Nikon Intensilight C-HGFIE Precentered Fiber Illuminator,Plan-Apo 100×/1.4 oil objective. Cells were cultured in YE medium at 28°C to late log phase, spotted on SPA and incubated at 28°C for 10–11 h. Zygotes suspended in EMM-N were sonicated briefly and mounted on a glass-bottom dish (Bioland) coated with 500 μg/mL lectin (Sigma). Images were acquired by Z sections and create Extended Depth of Focused (EDF) document. Image J software (NIH) was used to adjust brightness and contrast.

### Synchronous meiosis in *pat1-114* haploid cells

*pat1-114* haploid cells were used and induced to undergo synchronous meiosis as described previously(Yokobayashi and Watanabe 2005). Mid-log phase cultures were spun down at 2000 g for 5 min. The pelleted cells were resuspended in EMM-N and washed twice before being resuspended in EMM-N at a density of 0.2 OD660. Cultures were incubated for 6–16 h at 25°C to arrest cells in G1 phase. To induce synchronized meiosis, the culture was shifted to 34°C along with adding 1/50 volume of EMM. Aliquots of cells were collected at the indicated intervals. To monitor meiotic progression, aliquots of cells were stained with DAPI to observe the nuclear divisions.

### Protein preparation and immunoblot analysis

Cells (about 1 ×10^8^) were resuspended in 100 μL RIPA buffer (Beyotime, P0013B) and boiled for 5 min. About 1 mL of glass beads (about 500 µm in diameter) were added and the cells were broken by FastPrep-24 (MP Biomedicals) with 6.5m/s for 80s. A total of 10 µL of lysate from each sample was subjected to electrophoresis in 12% SDS-polyacrylamide gels and immunoblotted. Sgo1-GFP was detected with mouse anti-GFP monoclonal antibody (Transgene, Cat: HT801-01) at 1:5,000 dilution. GFP-PK-Moa1 was detected with mouse anti-PK monoclonal antibody (BioRad, Cat: MCA1360GA) at 1:5,000 dilution. Tubulin was detected with mouse anti-tubulin antibody TAT-1 (a gift from K. Gull) at 1:5,000 dilution. HRP-conjugated goat anti-mouse IgG (Transgene, Cat: HS201-01) was used as the secondary antibody at 1:5,000 dilution. Bands were visualized by using Clarity ECL substrate (RioRad, Cat: 170-5060) and captured using iBright 1500 imaging system (Invitrogen).

### Two-hybrid assays

The yeast two-hybrid system was used according to Clontech’s instructions (Cat: 630489). We amplified the ORFs of the *par1*^+^ gene by PCR and cloned it into pGBK-T7, a Gal4 DNA-binding domain–based bait vector. We also cloned *rec8*^*+*^, *rec8-2A* and *rec8-2E* into pGAD-T7, a Gal4 activation domain–based prey vector. Y2HGold and Y187 strains were used for transformation of pGBK-T7 and pGAD-T7, respectively. SD-tryptophan and SD-leucine plates were used as selective medium. Positive transformants of Y2HGold and Y187 were mixed on a YPDA plate to get diploid cells and streaked on SD-Trp-Leu plate to get single colonies; growth was checked on nutrition-restricted plates (SD-Trp-Leu-His, SD-Trp-Leu-Ade and SD-Trp-Leu-His-Ade).

### Statistical analysis

Data were analyzed using GraphPad Prism version 9.5.1 (GraphPad Software). To test for significant differences, we used one-way ANOVA with post-hoc Tukey’s multiple comparison test (Fig. 4E and 5A).

## Acknowledgements

We thank Silke Hauf for critically reading the manuscript, Jian Chen and Jingwen Zhou for general support, and the Yeast Genetic Resource Center (YGRC) for yeast strains and plasmids. We also thank Yuhei Goto for initial analyses of non-degradable Sgo1 and Moa1. This work was supported by the National Key Research and Development Program of China (2017YFC1600403), the National Natural Science Foundation of China (Key Program, 31830068), the National Key Research and Development Program of China (2023YFF1103701) and the Entrepreneurial and innovative talent of Jiangsu (JSSCRC2021495).

## Author contributions

Yongxin Liu: data curation, methodology, formal analysis, and investigation. Ke Zhang: investigation, methodology, formal analysis. Ziteng Bai: formal analysis. Li Sun: supervision, methodology. Haitong Hou: validation, funding acquisition. Yoshinori Watanabe: conceptualization, data curation, formal analysis, supervision, funding acquisition, validation, investigation, methodology, and writing—original draft.

---

**Figure EV1. Related to Fig. 3.**
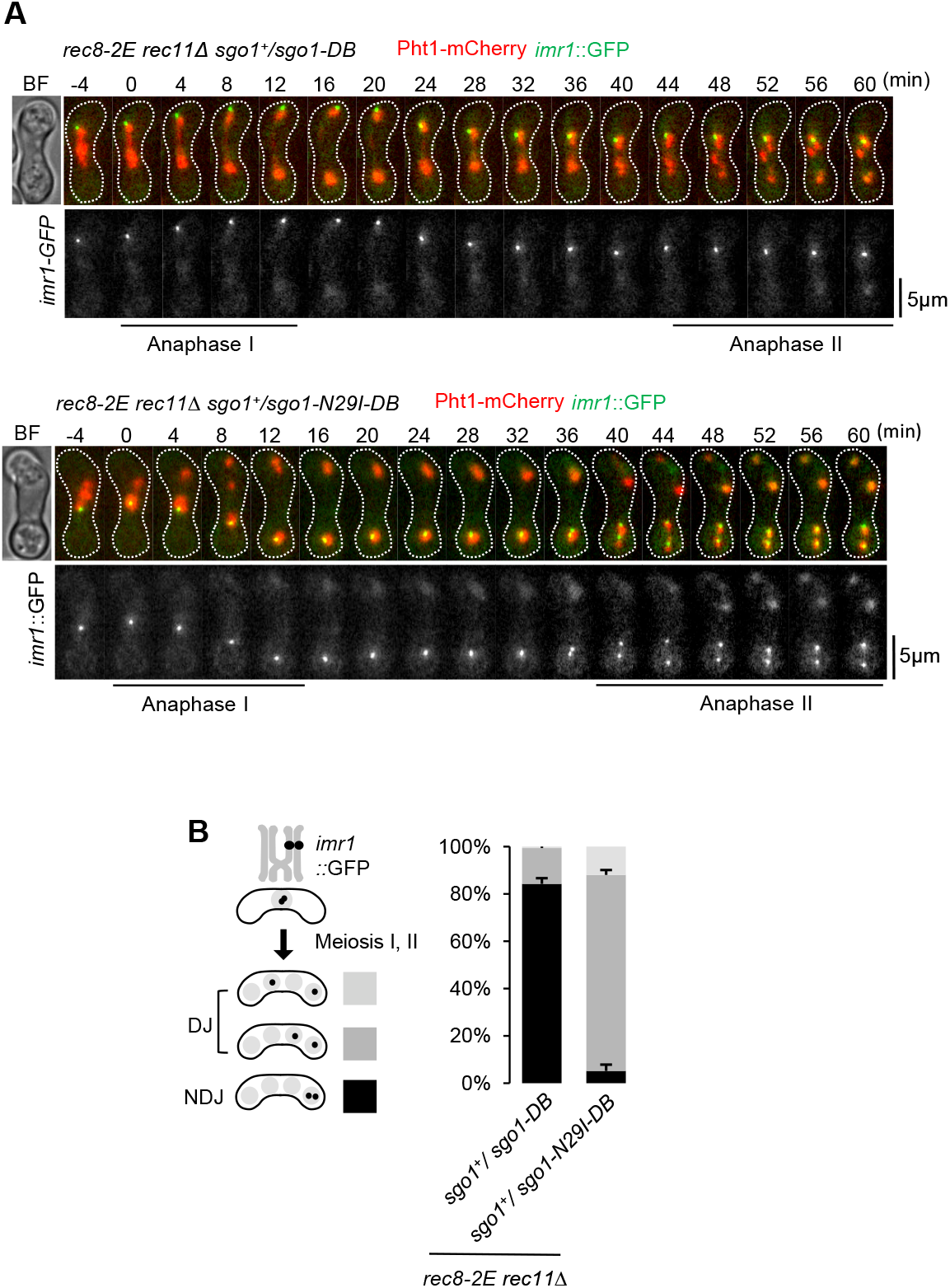
**A**, To complement the Fig. 3 experiments, we used *sgo1*^*+*^*/sgo1-N29I-DB* to measure the protection defects only in meiosis II because *sgo1-N29I-DB*/*sgo1-N29I-DB* may lose cohesion protection in meiosis I. As a control, we used comparable *sgo1*^*+*^*/sgo1-DB*. The representative examples of these strains are shown from the beginning of the first anaphase to the end of the second anaphase. Anaphase I and II are indicated by lines. Sister centromere behavior was observed in the indicated strains having *imr1*::GFP on only one homolog. DNA was visualized by expressing Pht1-mCherry from the endogenous locus. **B**, Segregation pattern of *imr1*::GFP marked on one homolog was monitored in the indicated cells after the meiosis II division.

**Figure EV2. Related to Fig. 6.**
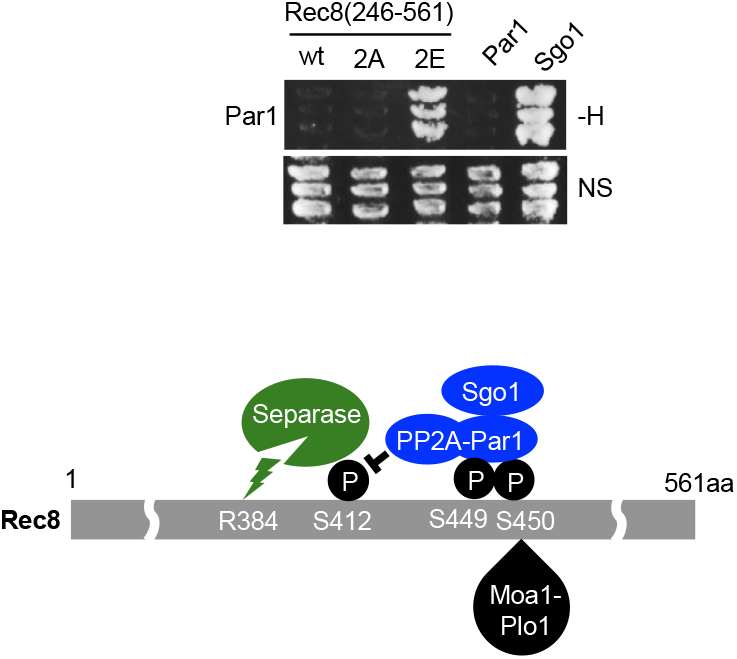
Two-hybrid assay indicates that Par1 interacts with Rec8-2E but not Rec8-WT or Rec8-2A. Schematic diagram of how Rec8 phosphorylation regulates Sgo1 and PP2A-Par1 localization and Rec8 cleavage by separase.

